# Developmentally-controlled generation of tRNA halves facilitates translational repression during sexual reproduction

**DOI:** 10.1101/2025.04.15.648877

**Authors:** Vasti Thamara Juarez-Gonzalez, Joan Marquez-Molins, Jinping Cheng, Juan Luis Reig-Valiente, Konstantin Kutashev, Stefanie Rosa, German Martinez

## Abstract

The role of transfer RNAs (tRNAs) as mediators between the genetic code and protein synthesis is well established. In parallel, tRNAs can generate different types of functional small RNAs (sRNAs) that accumulate during stress and certain developmental processes across diverse organisms. Interestingly, this class of sRNAs (termed tRNA-derived sRNAs, tsRNAs) accumulate in the male gamete of mammals, insects, and plants but their role in these reproductive cells is unclear. Here, we determine the molecular pathway of tsRNA biogenesis in the plant male gamete-containing structure (the pollen grain) and identify their role in mediating the characteristic translational repression taking place in this tissue. Our data demonstrates that male-accumulating tsRNAs are generated by a cell-controlled pathway to aid in mediating reproductive translational repression and that their accumulation is key to ensuring the proper function of the pollen grain.

## Introduction

tRNAs play a central role in translating genomic information into proteins, making them essential for cell viability^1^. Beyond this fundamental task, studies in multiple organisms have revealed that tRNAs possess a remarkable versatility, allowing them to regulate several cellular mechanisms due to their remarkable versatility. In particular, tRNAs can integrate environmental signals by generating different classes of tsRNAs,^1–5^ including tRNA-halves (28-36 nt) and shorter tRNA-derived fragments (tRFs, 18-28 nt)^6^. The accumulation of tsRNAs occurs under different stress conditions^2,7–9^ and, intriguingly, during gamete maturation in mammals, insects, and plants^10–13^.

Eukaryotic gamete development is a tightly regulated process involving a surprisingly evolutionary conserved transcriptional and epigenetic reprogramming^14–19^. In plants, the structure containing the male gametes, termed male gametophyte or pollen grain, undergoes a specific developmental program (Sup. Fig1a). This includes a series of mitotic divisions that take place after meiosis, resulting in a mature trinucleate cell containing two gametic or sperm cells (SCs) enclosed within a companion/nurse cell termed the vegetative cell (VC) which has its own nucleus (vegetative nucleus, VN). While critical for pollen tube development, the VC does not contribute genomic information to the next generation^15^. Prior to fertilization, the mature pollen grain (MPG) remains quiescent and developmentally arrested^20^. Similar to its analogous structure in animals, the sperm, the pollen grain undergoes transient translational repression during this quiescent stage accumulating most of the mRNAs required for fertilization^21,22^. While this repression may involve protein-mRNA interactions^23,24^, the molecular mechanisms orchestrating this process remain unknown.

Here, we have investigated the role of tsRNAs accumulating in the *Arabidopsis thaliana* MPG. Our data indicates that tRNA halves are present at high levels during male gamete maturation and that this accumulation correlates with a reprogramming of RNA modifications and increased expression of the Arabidopsis class II RNase T2, RNS2. Analysis of co-translational RNA degradation in the mature pollen grain reveals strong translational repression with limited mRNA degradation linked to tRNA halve accumulation. Furthermore, we found that tRNA halve depletion from the pollen grain inhibits its germination, indicating that their presence is essential for proper fertilization. In summary, we provide evidence that tRNA halves accumulation is a developmentally regulated process that naturally facilitates translation repression during Arabidopsis reproduction.

## Results

### tRNA-halves accumulate to high levels during male gamete development

We previously reported the accumulation of tRNA-derived fragments (termed tRFs) of 19-nts in the MPG of Arabidopsis and other plants where they posttranscriptionally regulate transposable elements^10^. Our initial study was technically limited to fragments smaller than 28-nts, prompting us to explore whether other types of tsRNAs also accumulated in this tissue. Northern blot analysis of different Arabidopsis tissues revealed that, together with tRFs, tRNA halves accumulate at high levels in the MPG compared to somatic tissues such as leaves or inflorescences (Fig 1a). Further analysis of tRNA halve accumulation by northern blot in different stages of pollen microgametogenesis showed that tRNA halve overaccumulation was not restricted to the mature gametophyte but was also detected in precursor stages, including uninuclear/microspore (UNM), binuclear (BNP), and trinuclear pollen (TNP) (Fig 1b). To characterize the composition of pollen-specific tRNA halves, we performed high-throughput sequencing of gel-purified sRNAs from leaves, UNM, BNP, TNP, and MPG and reanalyzed sRNA sequencing data from meiocytes^25^, the premeiotic cell that gives rise to 4 UNMs (Sup Fig 1a). This sRNA sequencing largely confirmed our northern blot detection, showing that all the stages of pollen grain development experience a high accumulation of tRNA halves (Fig 1c) that exhibit a characteristic size range of 32-36-nts, with a preference for 33-nt fragments in all the tissues analyzed (Fig 1c).

**Figure 1.**
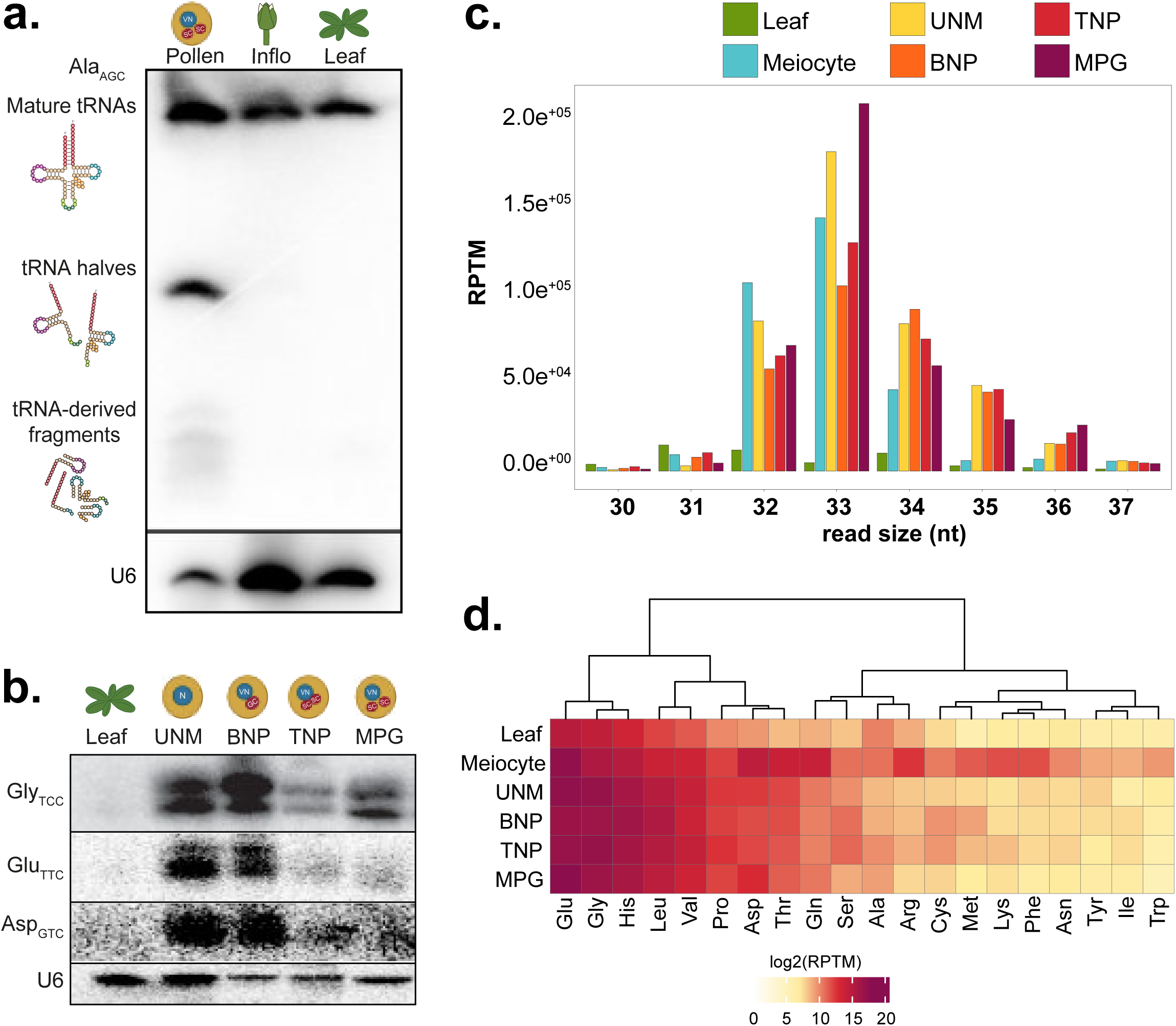
tRNA halves accumulate during Arabidopsis pollen development. **a.** Northern blot of total RNA from inflorescence, leaf, and mature pollen grain (MPG) using probes against 5′ tRNA halves derived from Ala-AGC and U6 as a loading control. **b.** Temporal profiling of tRNA halves by northern blot in isolated stages of pollen development: uninucleate microspore (UNM), binuclear pollen (BNP), trinuclear pollen (TNP), and MPG using probes for 5’tRNA halves from Glu-TTC, Gly-TCC, and Asp-GTC, and U6 as a loading control. **c.** Bar plot showing the size distribution of tRNA-mapped reads from small RNA sequencing (sRNA-seq) of leaf, meiocyte, UNM, BNP, TNP, and MPG. Reads are normalized by reads per ten million (RPTM) **d.** Heatmap of normalized (RPTM) abundance of tRNA-mapped reads separated by tRNA identity across somatic and reproductive tissues.

We sought to determine whether the accumulation of size-specific tRNA halves was consistent across different tRNAs, since the accumulation of specific tRNA halves has been previously reported in mouse sperm development and other organisms under stress conditions^26–28^. Similarly, all male reproductive maturation stages showed a similar accumulation profile, with halves derived from the Glu tRNA accumulating at the highest levels, followed by Gly, His, Leu, Val, Pro, and Asp (Fig 1d). Notably, the 32-33 nt Glu-TTC fragments were the most abundant in reproductive tissues compared to leaves, with the highest levels in MPG (Sup Fig 2a-b). To assess the conservation of this phenomena across eukaryotes, we analyzed public sRNA datasets from male gametes/gametophytes of diverse organisms, including insects (*D.melanogaster*), nematodes (*C.elegans*), mammals (*M.musculus*, *H.sapiens*), ferns (*M.polymorpha*), and plants from the monocotyledonous (*Z.mays*, *O.sativa*), and dycotiledoneous (*V.vinifera*, *C.rubella*) group. All these organisms exhibited a characteristic accumulation of 31-38 nt tRNA halves in their male gametes or gametophytes, indicating the evolutionary conservation of male gamete-associated tRNA halves (Sup Fig 2c). In summary, the Arabidopsis MPG displays a characteristic and specific accumulation of 33-nt tRNA halves, a feature conserved across multiple eukaryotic mature male reproductive tissues and/or cells.

### Pollen-specific tRNA halves are generated through a developmentally controlled pathway

The accumulation of pollen-specific tRNA halves in Arabidopsis led us to investigate the molecular events underlying their accumulation. In mammals, the generation of tRNA fragments has been linked to both a reduction in RNA modifications and increased activity of the RNase A angiogenin^7,28–33^, while in yeast, it has been associated with the activity of RNY1, a RNase of the T2 family^34^. To explore potential pathways governing tRNA halves biogenesis in the male gametophyte, we performed RNA sequencing from UNM, MPG, and leaves (as a somatic tissue control). Principal Component Analysis revealed distinct transcriptional profiles, with clear separation of pollen samples (UNM and MPG) and leaves. Additionally, UNM and MPG clustered separately, indicating overall transcriptomic differences during pollen development (Sup Fig 3a). Differential gene expression analysis of male reproductive tissues (UNM and MPG) compared to leaves revealed extensive transcriptional changes during pollen maturation, with 6,759/5,234 and 7,511/6,827 genes being upregulated/downregulated in the UNM and MPG, respectively (Fig 2a, Sup Fig 3b). Notably, although the UNM and MPG shared 50% of upregulated genes (Sup Fig 3c) the UNM exhibited a stronger enrichment in processes related to active cell proliferation while the MPG functions pointed to a transition from proliferative to specialized functions characteristic of MPG (Sup Fig 4).

**Figure 2.**
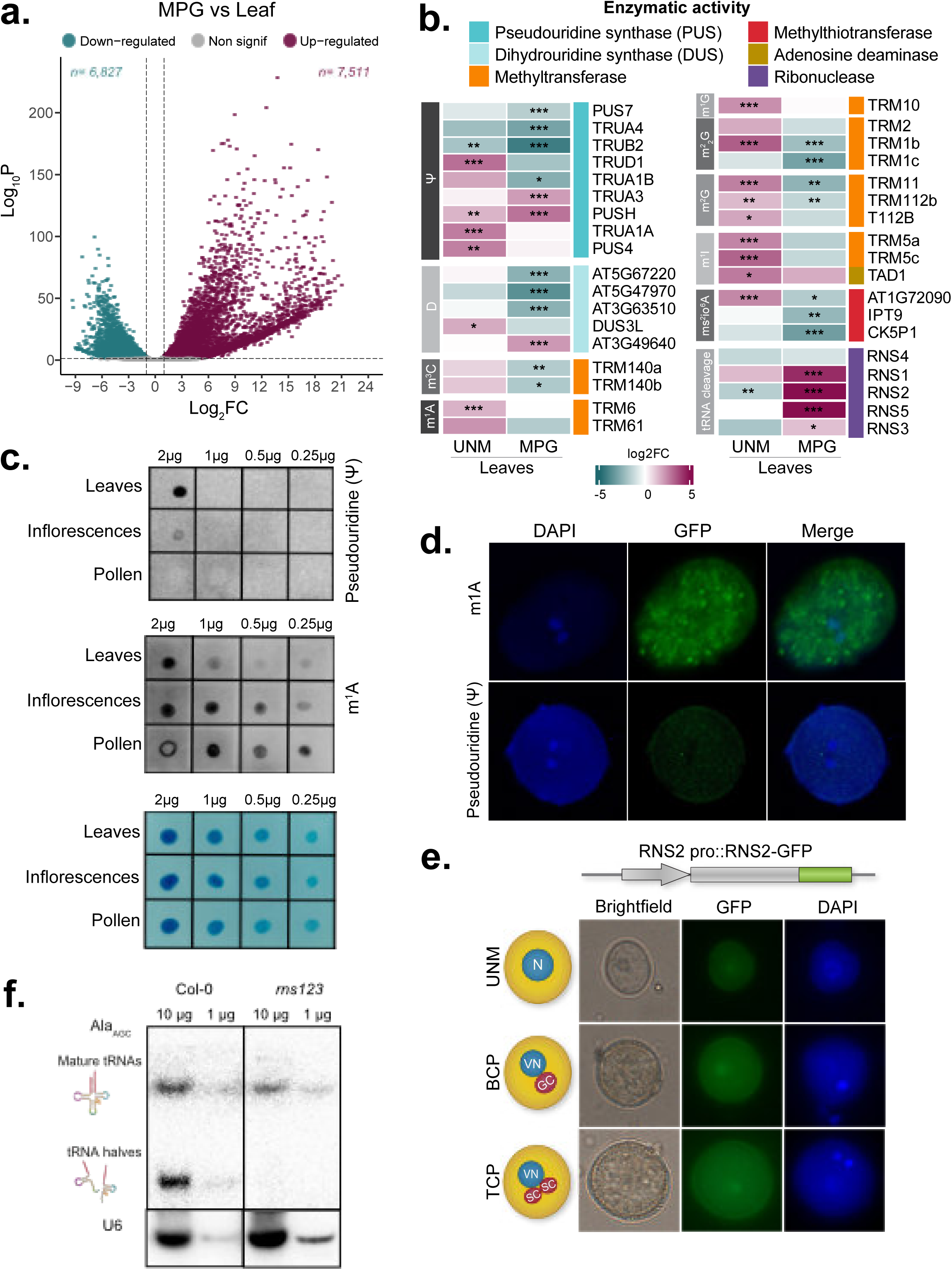
A developmentally controlled pathway regulates tRNA halves biogenesis in Arabidopsis pollen. **a.** Volcano plots showing differentially expressed (DE) genes in mature pollen grain (MPG) compared to leaves. Genes with adjusted p-value < 0.05 and log_2_Fold Change (log_2_FC) >= 1 or <= –1 were considered significantly upregulated (pink) or downregulated (blue), respectively. **b.** Heatmap of DE genes associated with enzymes that catalyze RNA modifications and RNase T2 enzymes in *Arabidopsis thaliana.* Comparisons between pollen and leaf tissues are shown (uninucleate microspore (UNM) vs. leaves and mature pollen grain (MPG) vs. leaves). RNA modification-related genes are grouped by enzymatic activity. **c.** RNA dot blot showing pseudouridine (Ψ) and N1-methyladenosine (m¹A) signals across Arabidopsis tissues. RNA loading ranges from 2 µg to 0.25 µg. **d.** Immunofluorescence detection of pseudouridine (Ψ) and N1-methyladenosine (m1A) in the MPG. Both images were obtained using similar exposure values. **e.** Subcellular localization of RNS2-GFP during different pollen maturation stages: uninucleate microspore (UNM), binuclear pollen (BNP), trinuclear pollen (TNP). **f.** Northern blot of total RNA (1 and 10ug) from Col-0 and *rns123* inflorescences using probes against 5′ tRNA halves derived from Ala-AGC and U6 as a loading control.

We then focused on genes previously reported to be involved in the biogenesis of tsRNAs in other species^35–39^, including those regulating the catalysis of different types of RNA modifications and Arabidopsis RNase T2 homologs (Fig 2b). tRNAs are among the most extensively modified RNA molecules (Sup Fig 3d) and changes in the presence of those modifications have been linked to the biogenesis of tsRNAs^7,30^. We identified that most genes involved in the RNA modification homeostasis (53%) were significantly downregulated in the MPG (Fig 2b). To confirm our RNA sequencing data, we generated translational fusions of several methyltransferases with high expression in the UNM or MPG, mediating m^1^A (TRM61), m^2^G (TRM112b), dihydrouridine (DUS1), and m^3^C (TRM140a). All except TRM140a showed strong accumulation in UNM but not in the MPG (Sup Fig 3e). From the extensive array of RNA modifications, m^1^A and pseudouridine (Ψ) are directly linked to tRNA stability^40,41^. To confirm that reduced activity of the m^1^A and Ψ catalysis pathways led to changes in the accumulation of these modifications, we compared their presence in MPG and somatic tissues by RNA dot blot (Fig 2c). Our results indicated major reprogramming of RNA modifications in the MPG, characterized by decreased Ψ and increased m^1^A levels (Fig 2c). This result was further corroborated by immunofluorescent detection of both modifications in the MPG using confocal microscopy (Fig 2d). Our findings align with previous genetic studies demonstrating that m^1^A is essential for gamete viability and the stability of the initiator methionyl-tRNA^42^. Indeed, we confirmed that mutants in the m^1^A methyltransferase TRM6 have a distorted segregation of their mutant allele when used as a pollen donor, indicating that this modification is key for pollen grain germination and/or fertilization (Sup Fig 3f).

tsRNA biogenesis has also been connected to increased activity of members of the RNase T2 family^34,37^. Functional RNases T2 possess a characteristic RNase T2 domain, containing two highly conserved active site motifs, CAS I and CAS II, which include histidine residues essential for catalytic ribonuclease activity^43^. The Arabidopsis RNase T2 family consists of five members (RNS1-5) that retain all the functional domains, with RNS2 being the closest homolog to RNY1 (Sup Fig 5a). An analysis of these genes in our RNA sequencing data, showed an upregulation of most of these genes in the MPG (Fig 2b). In addition, reanalysis of transcriptomic and proteomic data revealed that RNS2, was specifically upregulated in the MPG (Sup Fig 3g). To determine the subcellular localization of RNS2 activity, we generated a translational fusion of RNS2 to GFP (Fig 2e). Similar to angiogenin subcellular localization during stress^44^, RNS2 exhibited nucleo-cytoplasmic localization, suggesting it had access to both immature and mature tRNAs in the nucleus and cytoplasm (Fig 2e). To confirm the tsRNA generating activity of RNS genes, we generated an RNS1, RNS2, and RNS3 triple mutant. This triple mutant line displayed a diminished accumulation of tRNA halves (Fig 2f). Overall, our findings suggest that the coordinated activity of RNS2 and a decrease in the expression of genes involved in the catalysis of RNA modifications contribute to a developmentally controlled pathway that governs the biogenesis of tRNA halves during pollen development.

### 5’P-sequencing indicates that the pollen grain experiences extensive translational repression with reduced RNA degradation

tRNA halves have been implicated in mediating translational repression across multiple species^31,45,46^. To understand if our observed MPG-accumulating tRNA halves also had this effect, we performed a luciferase-based *in vitro* translation assay using synthetic 5’ tRNA halves. Fragments derived from Glu, Gly, Ala, and Asp tRNAs significantly reduced luciferase activity in a concentration-dependent manner (Fig 3a left panel and Sup Fig 6a). To determine whether endogenous RNAs also had this effect, we tested if total RNA and purified sRNAs fractions from the MPG had a similar effect over luciferase translation. Notably, both MPG-derived total RNA and, especially, sRNA purified-fraction inhibited luciferase activity (Fig 3a, right panel). Indeed, the inhibition mediated by sRNAs was similar to that caused by synthetic tRNA halves (Fig 3a, right panel). To understand the genome-wide mRNA dynamics in the male gametophyte of Arabidopsis, we performed 5’P-sequencing (5’P-seq) on the MPG and leaves. 5’P-seq allows the capturing of 5’monophosphate decay intermediates which are generated co-translationally and track the last translating ribosome, revealing ribosome dynamics at nucleotide resolution^47–49^. As such, a common characteristic of 5’P-seq datasets in multiple species is the accumulation of reads at 16/17 nts upstream the stop codon which indicate the slow translation termination step^50,51^. As expected, our 5’P-seq analysis in leaves revealed a profile showing this characteristic -16-nts upstream peak (Fig 3b). In contrast our 5’P-seq from the MPG exhibited a markedly different profile with overall less abundance of mRNA-mapped reads and the lack of the characteristic -16-nts termination peak (Fig 3b). This finding was reinforced when codon presence in the vicinities of the stop codon was compared between leaves and MPG (Fig 3c). While leaves showed an enrichment of 5′P signal specifically at - 16□nt upstream of termination codons, this was completely lost in the MPG, with no specific pausing signatures (Fig 3c and Sup Fig 6d and 6e). 5’P-seq data can be used to classify mRNAs according to their termination pausing as high or low^51^ (Fig 3d). While this classification displayed clear groups in our 5’P-seq libraries originating from leaves, with differential enrichment in functional categories (Sup Fig 7), using the same parameters for our MPG libraries identified a lower number of genes in both categories (69 and 66 in MPG against 100 and 127 in leaves) and failed to show clear termination peaks, with degradation appearing scattered and globally reduced (Fig 3d). Interestingly, although total 5’P-seq reads were lower in the MPG, we observed a relative enrichment of 5’P reads in the 5’UTR regions (Sup Fig 6b). In addition, reanalysis of poly (A) tail length data revealed that transcripts in the MPG, particularly those with high or low termination pausing, retained significantly longer poly (A) tails (Fig 3d, Sup Fig 6c) indicating an increased protection by poly (A)-binding proteins. Overall, our analysis of 5’P-seq in the MPG indicated that pollen transcripts have characteristics suggesting that they are translationally repressed and protected from degradation rather than being actively degraded^52^.

**Figure 3.**
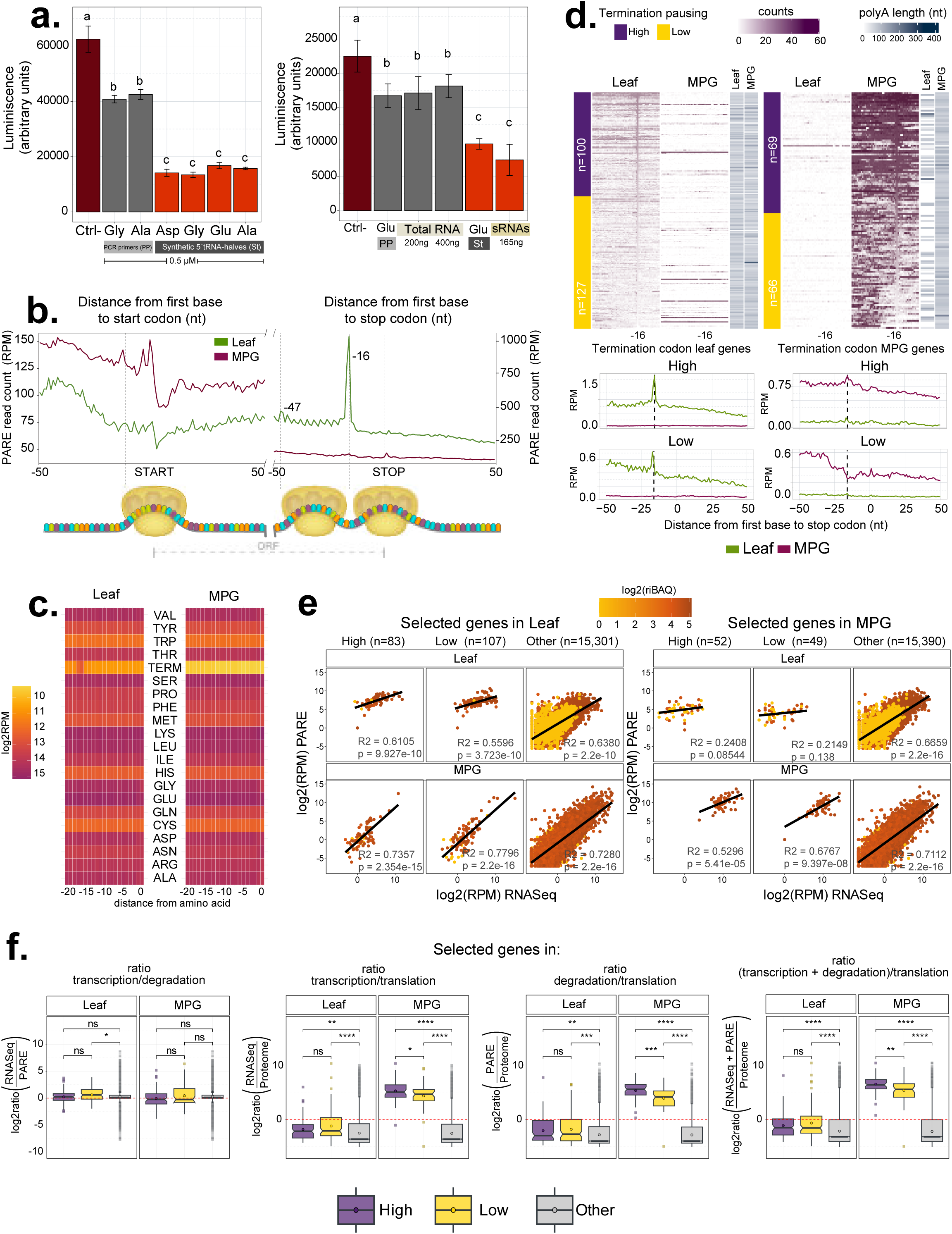
Translational repression and co-translational RNA decay dynamics. **a.** *In vitro* luciferase-based translation assay using synthetic 5’ tRNA halves (Glu, Gly, Ala, and Asp) at 0.5 µM or total RNA and small RNA fractions from mature pollen grain (MPG). DNA oligonucleotides (PP) used for the tRNA-halves (St) *in vitro transcription* were included as a control. Statistical differences were calculated using a two-way ANOVA analysis and Tukey’s HSD test, P□<□0.05. Differences between groups are shown with lowercase letters. **b.** Metagene profile for 5’P read coverage relative to ORF start and stop codon for 5’Pseq reads in MPG and leaves. **c.** Heatmap for aminoacid-specific 5’P coverage in MPG and leaves. Reads were normalized by reads per ten million (RPTM) and displayed on a logarithmic scale. **d.** Gene 5’P read coverage relative to ORF stop codon and poly(A) tail length in MPG and leaves (top). As previously reported by Zhang & Pelechano^42^, genes are sorted by the relative ribosome occupancy at the stop codon (-16 nt). Only genes with at least 10 reads per million (RPM) in this region were included and classified as a high or low termination codon (see methods). Metagene profile for 5’P read coverage relative to ORF stop codon for 5’Pseq reads in MPG and leave high and low termination pausing genes (bottom). **e.** Correlation between transcript (RNA-seq), RNA decay (PARE), and protein (riBAQ) levels in selected high, low, or non-defined (Other) pausing gene groups in Leaf and MPG. The statistical difference between tissues was calculated using a t-test, and the significance is shown below. ns: p-value >0.5; *:p-value<=0.5; ** :p-value<=0.1 and ***:p-value<=0.01 **f.** Ratios of transcription/degradation, transcription/translation, degradation/translation, and (transcription + degradation)/translation for gene sets with high, low, or non-defined pausing (Other) in Leaf and MPG. All the values correspond to MPG RNASeq, 5’Pseq and proteome. The statistical difference between pausing categories in each tissue was calculated using a t-test, and the significance is shown below. ns: p-value >0.5; *:p-value<=0.5; ** :p-value<=0.1 and ***:p-value<=0.01.

To further explore how transcription, RNA decay, and translation are coordinated in MPG and how these dynamics differ from leaves, we integrated our RNA-seq and 5’P-seq data with proteomic datasets from Arabidopsis leaves and pollen^22^. First, in both leaves and pollen, we observed a positive correlation between transcript abundance in RNA-seq and 5’P-seq (Fig 3e; Sup Fig 8a). Next, to better understand the balance between RNA metabolism and translation, we calculated transcription-to-degradation, transcription-to-translation, and degradation-to-translation ratios, along with a combined RNA dynamics-to-translation ratio ((transcription + degradation)/translation). These values describe how active a transcript is at the global RNA level compared to a protein product. Except for the transcription/degradation ratio (which remains similar across categories in MPG), all the other ratios were significantly higher for transcripts with high/low termination pausing in MPG and, moderately increased for the leaf-defined transcript sets (Fig 3f and Sup Fig 8b). Remarkably, the transcription/translation ratio was significantly higher in the MPG (Fig 3f, right panel) compared to leaves (Sup Fig 8b, right panel), indicating that in the MPG there is a reduced translation of highly transcribed mRNAs. In summary, our global analysis of transcription, co-translational decay, and translation indicates that in the MPG most mRNAs are actively produced but not translated. In addition, the characteristics of these mRNAs (longer poly(A)-tails and higher number of reads in the 5’ UTR) indicate that these might be protected from degradation or stalled in translation preinitiation complexes that could support gamete function after fertilization without new rounds of transcription.

### tRNA halves are important for proper pollen grain germination

The accumulation of tRNA halves in the MPG and their ability to repress translation *in vitro* suggested that they may play an important physiological role in pollen. Hence, we aimed to investigate the consequences of depleting specific pollen-enriched tRNA halves. To that aim we generated a short tandem target mimic (STTM) transgenic line, which block sRNA activity similar to a miRNA sponge^53–55^. We used this approach to target one of the most abundant tRNA halves in the pollen grain, Glu-TTC (Sup Fig 2a; Fig 4a). Northern blot analysis of the accumulation of Glu-TTC tRNA halves in these transgenic lines confirmed their ability to reduce the accumulation of this tsRNA (Fig 4a, right panel). Phenotypic analysis of this line and our previously generated *rns1/2/3* triple mutant, revealed that pollen germination was impaired in all lines with disrupted tRNA halves accumulation, with the Glu- STTM line showing the most severe reduction (Fig 4b-c). Notably, neither pollen viability nor vegetative development (rosette leaf number) was affected in these lines (Sup Fig 9a-b), indicating that the germination defects were not due to broader developmental abnormalities. Interestingly, functional enrichment of genes with high or low termination pausing in pollen was associated with pollen tube growth (Sup Fig 7), suggesting that translationally repressed mRNAs may be important for pollen germination and support the idea that tRNA halves contribute to posttranscriptional control by modulating the translation of those mRNAs.

**Figure 4.**
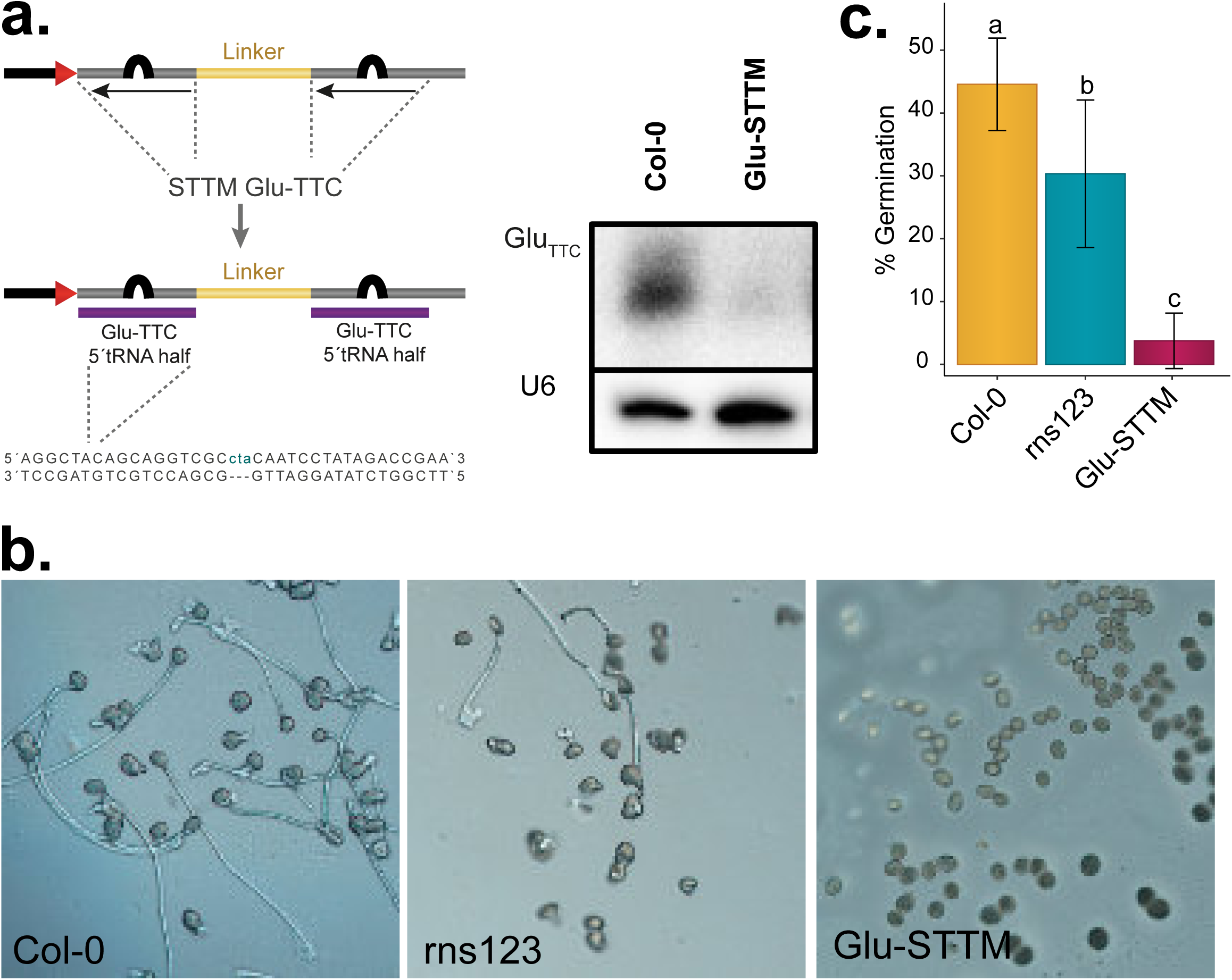
tRNA halves are required for proper pollen germination. **a.** Schematic representation of short tandem target mimic (STTM) constructs targeting the 5′ tRNA half derived from Glu-TTC. The STTM contains a central linker and two imperfect antisense binding sites that sequester the target tRNA half. The sequence of Glu-GTC 5′ half is shown below. The right panel shows a northern blot of total RNA from control and the Glu-STTM transgenic line using a probe against a 5′ tRNA halve derived from Glu-TTC and U6 as a loading control. **b.** Representative images of i*n vitro* pollen germination analysis in wild-type (Col-0), the rns123 triple mutant, and the STTM line targeting Glu-TTC (Glu-STTM) tRNA halves. Scale bars, 50□μm. **c.** Barplot showing pollen germination efficiency in wild-type (Col-0), rns123, and Glu-STTM lines. Analysis was performed using three biological replicates with three technical replicates each one. Error bars indicate standard deviation. Statistical differences were calculated using two-way ANOVA analysis and Tukey’s HSD test, P□<□0.05.

## Discussion

Sexual reproduction in eukaryotes is orchestrated by strong epigenetic and transcriptomic reprogramming, which are key to ensuring the success of the fertilization process. The plant male reproductive structure, the MPG, represents the culmination of plant development and the transition to the next generation. This structure is fundamental for the transfer of genomic information to the next generation and, as such, the molecular processes taking place in this tissue must be tightly regulated. Translational repression in the MPG of different plant species has been identified and partially studied^22–24,56^, although it was not well understood. Here, we have shed light into how that process is regulated by (1) identifying the natural accumulation of tRNA halves in the MPG, (2) describing the biogenesis pathway of these sRNAs, (3) analyzing their translational repression activity, and (4) determining their physiological role in the MPG.

tRNA halve accumulation is a well-known response to different stresses in multiple organisms^2,7–9^. In addition, several works have reported the accumulation of tsRNAs in reproductive tissues, especially in the sperm^10–13^. Our work describes in detail the characteristics of pollen-accumulating tsRNAs and the conservation of this accumulation in multiple male gametic tissues from a diverse range of eukaryotes (Sup Fig 2c). Furthermore, we describe the molecular pathway regulating the accumulation of pollen-derived tRNA halves, which is led by increased activity of the RNS2 and a developmentally-controlled reprogramming of RNA posttranscriptional modifications (Fig 2). Due to the lack of angiogenin homologs, RNS genes have been shown to mediate the biogenesis of different tsRNAs in Arabidopsis^37,57^. Our data indeed shows that RNS genes mediate tRNA halve biogenesis, in particular the pollen-expressed RNS2. This result also indicates a potential tissue-specific regulation of RNS activity and, subsequently, of tsRNA accumulation. Furthermore, we connect tRNA halve biogenesis to a reprogramming of RNA modifications. tsRNA biogenesis has been previously linked to decreased RNA modifications associated with different types of stress^7,30^. Interestingly, our data indicates that RNA modifications are reprogrammed in the male reproductive tissues of Arabidopsis which exhibits higher levels of m^1^A and reduced levels of Ψ (Figure 2). It is plausible to speculate that although a partial reprogramming of RNA modifications is needed, the role of m^1^A in mediating the stability of the initiator tRNA Met-CAT^42^ needs to be maintained. In addition, the role of Ψ in the maintenance of the stability of the tRNAs folded structure is well-known^58^, indicating that the reprogramming of this modification could lead to the accumulation of unmodified tRNAs that could be targets of RNS cleavage.

Our work also shows that pollen accumulating tRNA halves drive translational repression both in vitro and in vivo. Indeed, our use of 5’P-seq to track co-translational mRNA decay in the MPG (Fig 3b) allowed us to confirm the extensive translational repression taking place in this structure (as previously described^21,24,52^) which happens in the absence of mRNA degradation (Fig 3a and d). Connected to that role of tRNA halves in the MPG, we confirmed that the tRNA halve-mediated translational repression is physiologically relevant for pollen development since direct removal of tRNA halves using genetic mutants (*rns123*) or an STTM transgenic line, removing one of the highest accumulating tRNA halves (Glu-TTC), compromise pollen grain germination (Fig 4c).

We hypothesize that, since reproductive tissues suffer reduced Pol II activity^21,22^, tRNA halves dependence on Pol III might make them an extensive reservoir of sRNAs. In addition, tRNA halves might represent a highly unspecific/general manner to regulate mRNA translation since they have been reported to bind directly to the ribosome^59,60^, in opposition to other sRNAs with translational repression activity such as miRNAs^61^. There are nevertheless several open questions concerning the activity and role of such sRNAs in reproductive tissues. For example, it is unknown if tRNA halves interact with RNA binding proteins to promote their stabilization and facilitate their access to mRNAs. In addition, it remains to be proven if the biogenesis route described here for Arabidopsis is responsible for the biogenesis of sperm-accumulating tRNA halves in other organisms and/or under stress conditions. In summary, our findings demonstrate that developmentally regulated tsRNAs are extremely versatile molecules with significant physiological roles during plant sexual reproduction. In addition, our work identifies yet another sRNA-based regulatory mechanism playing important roles in sexual reproduction showing the importance of these molecules for regulating multiple aspects of this process.

## Methods

### Plant material

*Arabidopsis thaliana* plants were grown under standard long-day conditions (16-h light and 8-h dark) in a growth chamber illuminated with fluorescent tubes (Philips F25T8/TL841 25 watt tubes with a light intensity of 130 μmol m^−2^ s^−1^) at 22□°C. Plants were sown into potting soil (S-Jord, Hasselfors Garden). The mutant alleles used in this study were: *rns1* (GK- 760D11-024594), *rns2* (SALK_069588), and *rns3* (SALK_041269).

### Total RNA, sRNA RNA gel blot and Northern blot assays

Total RNA was isolated using TRIzol reagent (Life Technologies). For small RNA gel blot detection, 20 μg of total RNA were loaded in each lane for each pollen developmental stage, leaf and inflorescence sample. sRNA gel electrophoresis, blotting, and cross-linking were performed as described in Pall and Hamilton (2008)^62^. Membranes were hybridized in ULTRAhyb^®^-Oligo hybridization buffer (Thermo Fisher Scientific) with oligonucleotide probes targeting Ala-AGC, Glu-TTC and Asp-GTC 5’tRNA half fragments, which were end-labeled with [γ-32P] ATP (PerkinElmer) using T4 polynucleotide kinase (New England Biolabs).

### RNA-seq, small RNA, and 5’P-seq library construction

sRNA libraries were constructed as described in McCue et al. (2012)^63^ adapted for Arabidopsis pollen. 5’P-seq libraries were prepared following the protocol from Carpentier et al. (2021)^64^ adapted to pollen tissues and using mRNA-enriched fractions obtained with the NEB Magnetic mRNA Isolation Kit (New England Biolabs). sRNA libraries were made using the NEBNext Small RNA Library Prep Set for Illumina (New England Biolabs) with gel-enriched sRNAs as described in Martinez et al. (2016)^65^. RNA-seq libraries were prepared using the NEBNext® Ultra™ II Directional RNA Library Prep Kit for Illumina® (New England Biolabs) and barcoded using the NEBNext® Multiplex Oligos for Illumina® kit (New England Biolabs). For sRNA, RNA-seq, and 5’P-seq libraries, three biological replicates were used per tissue or genotype, each consisting of pooled samples from 18-20 plants.

### Pollen grain separation, germination, viability test, and microscopy

Pollen grain fractionation was performed as described in Dupl’akova et al. (2016)^66^. The pollen developmental stages used for sRNA sequencing correspond to the fractions termed B1 (Uninuclear), B3 (Binuclear), and A3 (Trinuclear). Pollen germination was assessed using the media recipe from Rodriguez-Enriquez et al. (2013)^67^ with minor modifications. Germination assays were performed over three independent experiments, each with three technical replicates. After 24h ∼500 pollen grains per replicate were scored as germinated or non-germinated. Statistical analysis was performed using two-way ANOVA followed by Tukey’s HSD test (P□<□0.05). Pollen viability was assessed using Alexander staining (Alexander, 1969)^68^. Imaging of germinated pollen grains and stained grains was performed in a Leica DM RX microscope. For pollen viability quantification, 700-800 pollen grains per genotype were scored as viable or aborted. GFP fluorescence of RNS2-GFP pollen grains was visualized on slides containing 50% glycerol using either a Zeiss Axioplan or a Leica DMI 4000 microscope.

### In vitro luciferase translational assay

To synthesize tRNA halves, forward and reverse oligonucleotides containing the T7 promoter sequence upstream of the corresponding tRNA halve sequence were annealed by incubating them at 95°C for 10 min, followed by gradual cooling to 25°C (10°C every 2 min). Annealed oligos were used as templates for *in vitro* transcription, which was performed using MAXIscript™ T7 Transcription Kit (Thermo Fisher Scientific). For translational inhibition assays, in vitro luciferase translation was carried out using the TnT™ Coupled Wheat Germ Extract System (Promega), Synthetic tRNA halves were added at final concentrations of 0.2 and 0.5 ng per reaction. Luciferase activity was measured using a Fluostar Omega fluorometer (BMG Labeltech).

### Immunofluorescence detection

Immunofluorescence detection of RNA modifications in the MPG was performed as previously described^69^. In brief, opened flowers were collected in PBS containing 0.05% tween20 and centrifuged at 2000 g for 2 min to collect the MPG. Samples obtained from the collected MPG were fixed using 4% paraformaldehyde for 30□minutes at room temperature, washed 3 times with PBS buffer, and digested using a digestion buffer (0.4% Cytohelicase, 1% Polyvinylpyrollidone, 1.5% Sucrose) at 37□°C for 60□min. After this, the samples were washed with PBS for 3 times, treated with 0.1% Triton X-100 for 30□min, washed again 3 times with PBS, and blocked with 1% BSA at 37□°C for 1□h. After this treatment, samples were incubated with the corresponding antibodies (anti-m1A, Diagenode, C15200235; or anti-Pseudouridine, Diagenode, C15200247) overnight at 4□°C in 1% BSA. After this incubation the samples were washed 3 times with PBS and incubated with the secondary antibody anti-Alexa Fluor 488 at 37 °C for 1h. Finally, the samples were washed 3 times with PBS, incubated with DAPI for 10 minutes and subsequently analyzed using a LSM780 confocal microscope (Zeiss).

### Bioinformatic analyses

Protein Ids for RNases T2 homologs were inferred using Phytozome version 13^70^. Protein sequences were obtained using Ensembl Plants^71^. Functional domain analysis was performed using Pfam^72^, and only sequences containing a RNase T2 domain were retained. Multiple sequence alignments were conducted in MUSCLE^73^, and phylogenetic reconstructions were inferred in MEGA 11^74^, using the Maximum Likelihood method and Whelan And Goldman model^75^. A discrete Gamma distribution was used to model rate heterogeneity among sites. sRNA, RNA-seq, and 5’P-seq libraries were trimmed using Trim Galore v0.6.12^76^. Trimmed and size-selected sRNA reads were aligned to the Arabidopsis thaliana TAIR10 genome using bowtie (v1.1.2)^77^ allowing up to two mismatches (-v2). tRNA annotations were obtained from Genomic tRNA Database (GtRNAdb)^78^. Feature counts were obtained using HTseq v.0.11.1^79^, and expression levels were normalized as reads per ten million (RPTM) to the total reads mapped to the *A. thaliana* chromosomes. tRNA read counts where further normalized copy number per each amino acid isoaceptor arm identity.

RNA-seq reads were aligned to TAIR10 using STAR v2.7.11b^80^. Raw counts were normalized by RPTM. Differential gene expression and principal component analyses (PCA) were conducted in R v4.4.3^81^ using DESeq2 v1.44.0^82^. Volcano plots and heatmaps were generated with EnhancedVolcano v1.22.0^83^ and ComplexHeatmap v2.22.0^84^, respectively. Functional enrichment was performed using ShinyGO v0.82^85^ and summarized using REVIGO v1.8.1^86^. All other plots were generated with ggplot2 v3.5.1^87^.

Trimmed 5’P-seq reads were mapped using STAR 2.7.11b, and BAM files were analyzed with Fivepseq^50^ (http://pelechanolab.com/software/fivepseq), to assess 5’P ends relative to start, stop codons, and codon specific pausing. Reads uniquely mapped to protein-coding genes were normalized to reads per million (RPM), and metagene plots were constructed based on the summed counts at each relative. Only genes with >10 RPM in the -14 to -47 nt window upstream of the stop codon were considered for classification. Genes with a signal above or below the global mean were classified as a high or low termination pausing, respectively. Proteomic data for Arabidopsis pollen and leaf tissues were obtained from Mergner et al. (2020)^88^. Statistical analyses were conducted using tidyr and dplyr R packages^87^. Heatmaps of pausing categories were generated using a complex heatmap R package and boxplots, bar plots, and correlation plots were produced using ggplot2.

## Acknowledgments

We thank Formas (2021-01161), the Swedish Research Council (VR 2021-05023), and the Knut and Alice Wallenberg Foundation (KAW 2019.0062) for supporting research in the Martinez group. The data handling was enabled by resources provided by the Swedish National Infrastructure for Computing (SNIC) at UPPMAX partially funded by the Swedish Research Council through grant agreement no. 2018-05973.

---

**Supplementary Figure 1. *Arabidopsis thaliana* male gametogenesis and conservation of tRNA halves across species. a.** Diagram summarizing the key stages of male gametophyte development in *Arabidopsis thaliana*. Following meiosis, the haploid uninucleate microspore (UNM) undergoes a mitotic division to form the binuclear pollen grain (BNP) composed of a vegetative nucleus (VN) and a generative cell (GC). The GC undergoes a second mitosis to generate two sperm cells (SCs), resulting in the trinuclear pollen grain (TNP). This final structure, the mature pollen grain (MPG), contains one VN and two SCs in a shared cytoplasm. Upon pollination, the MPG germinates and extends a pollen tube toward the ovule. The VN promotes pollen tube growth, while the two SCs are transported to the female gametophyte. Double fertilization involves one SC fusing with the egg cell to produce the diploid zygote (2n) and the other SC fusing with the central cell to generate the triploid endosperm (3n), which nourishes the developing embryo. The resulting embryo and endosperm develop within the protective seed coat, forming a seed that will undergo maturation, followed by a desiccated dormant state. Germination occurs upon exposure to water and light, giving rise to a new flowering plant.

**Supplementary Figure 2. tRNA-specific enrichment of tRNA halves in Arabidopsis somatic and reproductive tissues. a.** Heatmap of the normalized abundance of tRNA-mapped reads grouped by tRNA codon identity in somatic and reproductive tissues. **b.** Heatmap of normalized abundance of tRNA-mapped reads grouped by codon and read size, only reads from 30 to 37 nt are shown. Data is normalized by reads per ten million (RPTM) and displayed on a logarithmic scale. **c.** Heatmap showing the size distribution profiles of tRNA mapped reads from sRNA-seq datasets of somatic and male reproductive tissues across multiple eukaryotic species, including *C. elegans, D. melanogaster, M. musculus, H. sapiens, M. polymorpha, Z. mays, O. sativa, V. vinifera*, and *C. rubella*.

**Supplementary Figure 3. Transcriptional analysis identifies a differential activity of RNA modification pathways in the MPG. a.** Principal component analysis (PCA) of RNA-seq data from leaf, uninucleate microspore (UNM), and mature pollen grain (MPG). **b.** Volcano plots showing differentially expressed (DE) genes in UNM compared to leaves. Genes with adjusted p-value < 0.05 and log_2_Fold Change (log_2_FC) >= 1 or <= –1 were considered significantly upregulated (pink) or downregulated (blue), respectively. **c.** Venn diagram showing overlap of significantly up and down-regulated genes between UNM and MPG. **d.** Schematic representation of the secondary tRNA structure highlighting major nucleotide modifications and their positions. **e.** Analysis of the subcellular accumulation of translational reporter fusions of selected RNA modification enzymes (TRM61, TRM112b, TRM140a, DUS1) in leaves, UNM and MPG. **f.** Analysis of the segregation of the mutant alleles trm6 and trm61 for the cross trm6/+ and trm61/+. **g.** Protein and transcript abundance of RNS gene family expression during pollen development obtained from the reanalysis of proteomic and transcriptomic datasets.

**Supplementary Figure 4. Functional enrichment of differentially expressed genes in pollen compared to leaves.** Dot plots showing Gene Ontology (GO) enrichment analysis for upregulated genes in UNM (left, in purple), MPG (right, in purple) compared to leaves or downregulated genes in UNM (left, in blue), MPG (right, in blue) compared to leaves. Enriched terms are grouped into Biological Process (BP), Molecular Function (MF), and Cellular Component (CC). The circle size indicates the percentage of genes enriched in each category, the color intensity reflects FDR significance, and the x-axis represents each term’s Fold Enrichment.

**Supplementary Figure 5. Conservation and domain organization of RNase T2 proteins.** RNase T2 family phylogenetic reconstruction using amino acid sequences from Arabidopsis (At), rice (Os), maize (Zm), *Vitis vinifera* (Vv), *Saccharomyces cerevisiae, Caenorhabditis elegans and Drosophila melanogaster.* T2 ribonuclease domain organization and divergence among RNS family members are also shown. Multiple sequence alignments of the CAS I and CAS II motifs from RNase T2 family proteins highly conserved histidine residues are highlighted. Phylogenetic Reconstruction was built using Molecular Evolutionary Genetics Analysis version 11, using the maximum verisimilitude method with a gamma-distributed model and 500 bootstraps.

**Supplementary Figure 6. Translational repression, poly (A) tail gene length, and co-translational RNA decay dynamics in MPG and leaves. a.** *In vitro* luciferase-based translation assay using synthetic 5’ tRNA halves (Glu, Gly, Ala, and Asp) at 0.2 µM. DNA oligonucleotides (PP) used for the tRNA-halves (St) *in vitro transcription* were included as a control. Statistical differences were calculated using a two-way ANOVA analysis with a Tukey HD post hoc test. Differences between groups are shown with lowercase letters. **b.** 5’P read coverage in 5′UTR, CDS, and 3′UTR regions in leaves and MPG. Reads were classified relative to ORF start and stop codon. **c.** Poly(A) tail length distributions of high, low, and other pausing gene groups. The statistical difference between tissues was calculated using a t-test, and the significance is shown below. ns: p-value >0.5; *:p- value<=0.5; ** :p-value<=0.1 and ***:p-value<=0.0. **d.** Heatmap for codon-specific 5’P coverage, including termination codons (TAA, TAG, TGA) in MPG and leaves. **e.** Heatmap for Dipeptide -specific 5’P in MPG and. Reads were normalized by reads per ten million (RPTM) and displayed on a logarithmic scale.

**Supplementary Figure 7. Functional enrichment of co-translationally degraded transcripts by termination pausing gene group.** Dot plots showing Gene Ontology (GO) enrichment analysis for high and low termination pausing in leaves and MPG. Enriched terms are grouped into Biological Process (BP), Molecular Function (MF), and Cellular Component (CC). The circle size indicates the percentage of genes enriched in each category, the color intensity reflects FDR significance, and the x-axis represents each term’s Fold Enrichment.

**Supplementary Figure 8. Transcription, translation, and co-translational RNA decay dynamics in leaf-selected categories. a.** Correlation between transcript (RNA-seq), RNA decay (PARE), and protein (riBAQ) levels in selected high, low, or non-defined (Other) pausing gene groups in leaf and MPG. The statistical difference between tissues was calculated using a t-test, and the significance is shown below. ns: p-value >0.5; *:p- value<=0.5; ** :p-value<=0.1 and ***:p-value<=0.01. **b.** Ratios of transcription/degradation, transcription/translation, degradation/translation, and (transcription + degradation)/translation for gene sets with high, low, or non-defined pausing (Other) in Leaf and MPG. All the values correspond to Leaf RNASeq, 5’Pseq and proteome. The statistical difference between pausing categories in each tissue was calculated using a t-test, and the significance is shown below. ns: p-value >0.5; *:p-value<=0.5; ** :p-value<=0.1 and ***:p-value<=0.01.

**Supplementary Figure 9. Pollen viability and vegetative development in STTM lines and rns123 mutants. a.** Representative images of Arabidopsis plants at bolting stage from wild-type (Col-0), the rns123 triple mutant (rns123), and the STTM line targeting Glu-TTC (Glu-STTM) tRNA halves (left). The violin plot shows the leaf number in all the lines (right) at this stage. The statistical difference between tissues was calculated using a t-test, and the significance is shown below. ns: p-value >0.5; *:p-value<=0.5; ** :p-value<=0.1 and ***:p- value<=0.01. **b.** Representative images showing pollen viability in wild-type (Col-0), rns123 and, Glu-STTM line using an Alexander staining technique (left). The table shows the number of viable and aborted pollen grains in the indicated lines (right, top). Bar plot showing the percentage of viable pollen grains in Col-0, rns123, and Glu-STTM (right, bottom). **c.** Principal Component Analysis (PCA) of RNA-seq data from mature pollen grains (MPG) of wild-type Col-0, rns123 triple mutant, and the STTM transgenic line targeting Glu- TTC 5′ tRNA halves. Each point represents one biological replicate. **d.** Volcano plots show differentially expressed genes (DE) between Col-0 and the rns123 mutant (top) and, Glu- STTM (bottom). Genes with adjusted p-value < 0.05 and log_2_Fold Change (log_2_FC) >= 1 or <= –1 were considered significantly upregulated (pink) or downregulated (blue), respectively. Genes with the biggest and most significant DE are annotated. **e.** Venn diagrams showing the overlap of upregulated (top) and downregulated (bottom) genes across all comparisons with Col-0.

