## Supplementary Figures for "Developmentally-controlled generation of tRNA halves facilitates translational repression during sexual reproduction"

#### Supplementary Figure 1

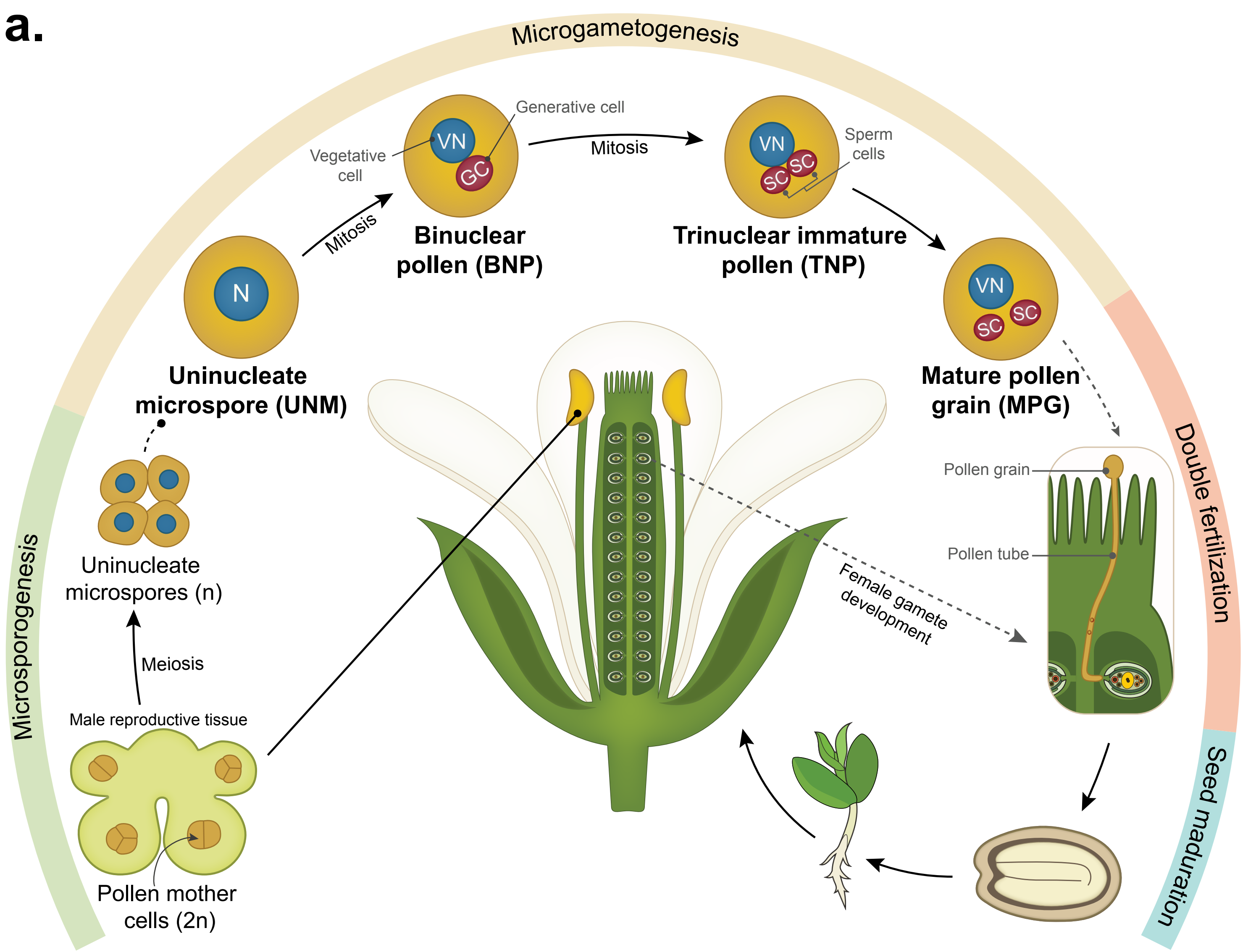

#### Supplementary Figure 2

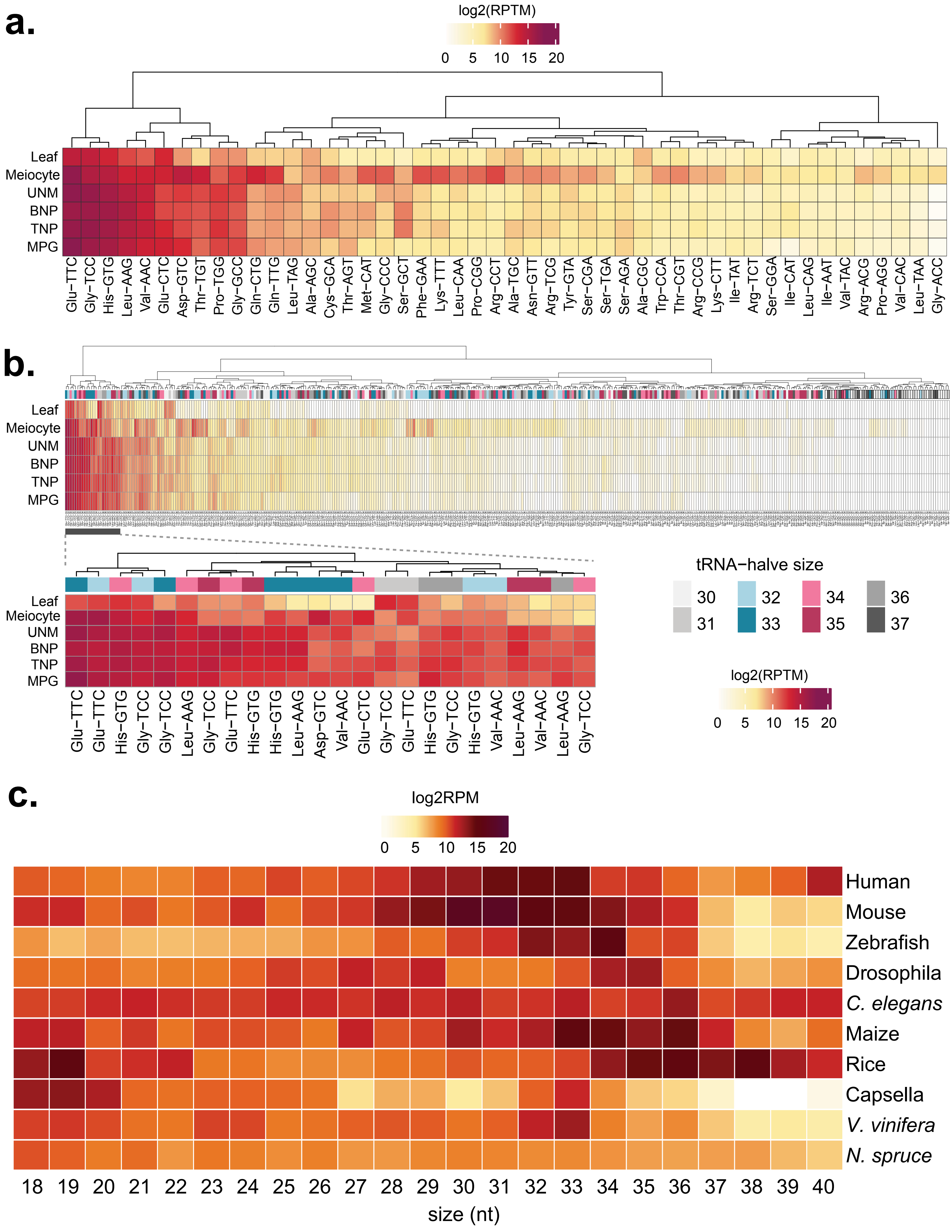

### Supplementary Figure 3

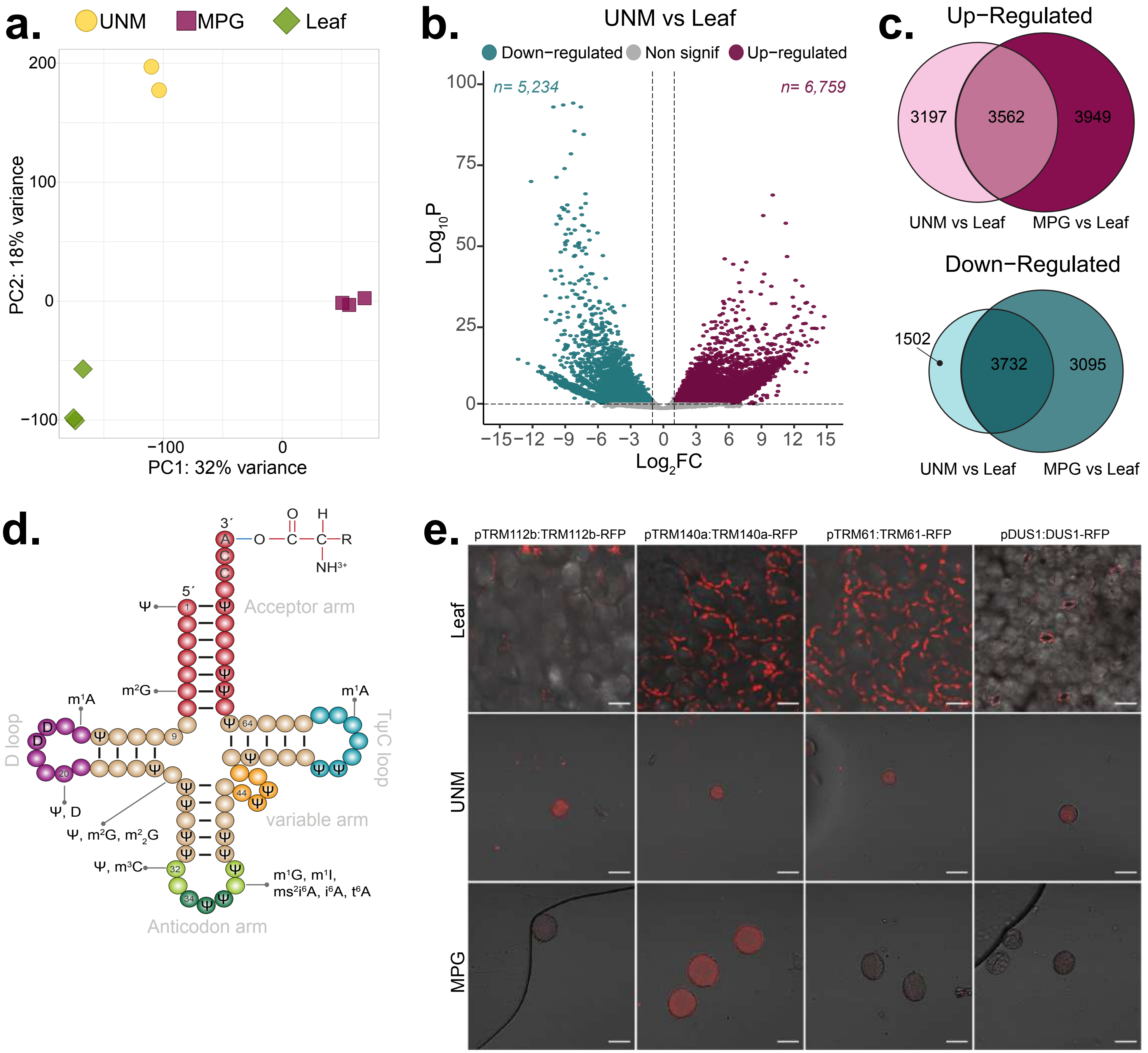


**f.**

***trm61-1/+* X *trm6/+***

| | Expected | Observed |
|---|---|---|
| *trm61/+* | 44/45 (50%) | 41 (46%) |
| *trm6/+* | 44/45 (50%) | 11 (12%) |
| *trm61/+ trm6/+* | 22/23 (25%) | 9 (10%) |

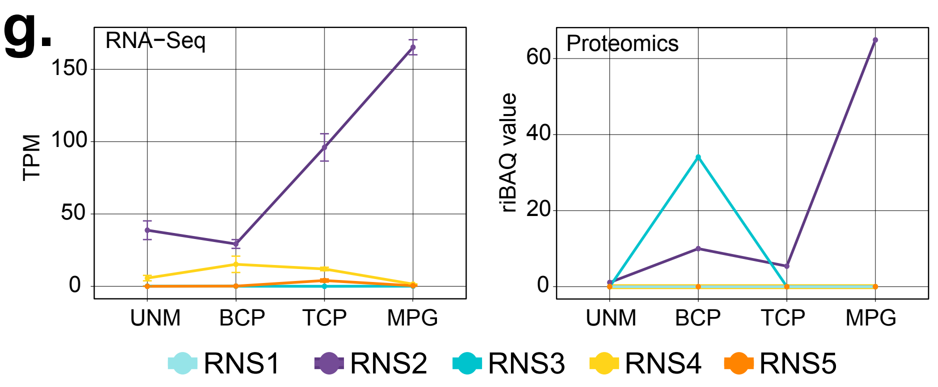

### Supplementary Figure 4

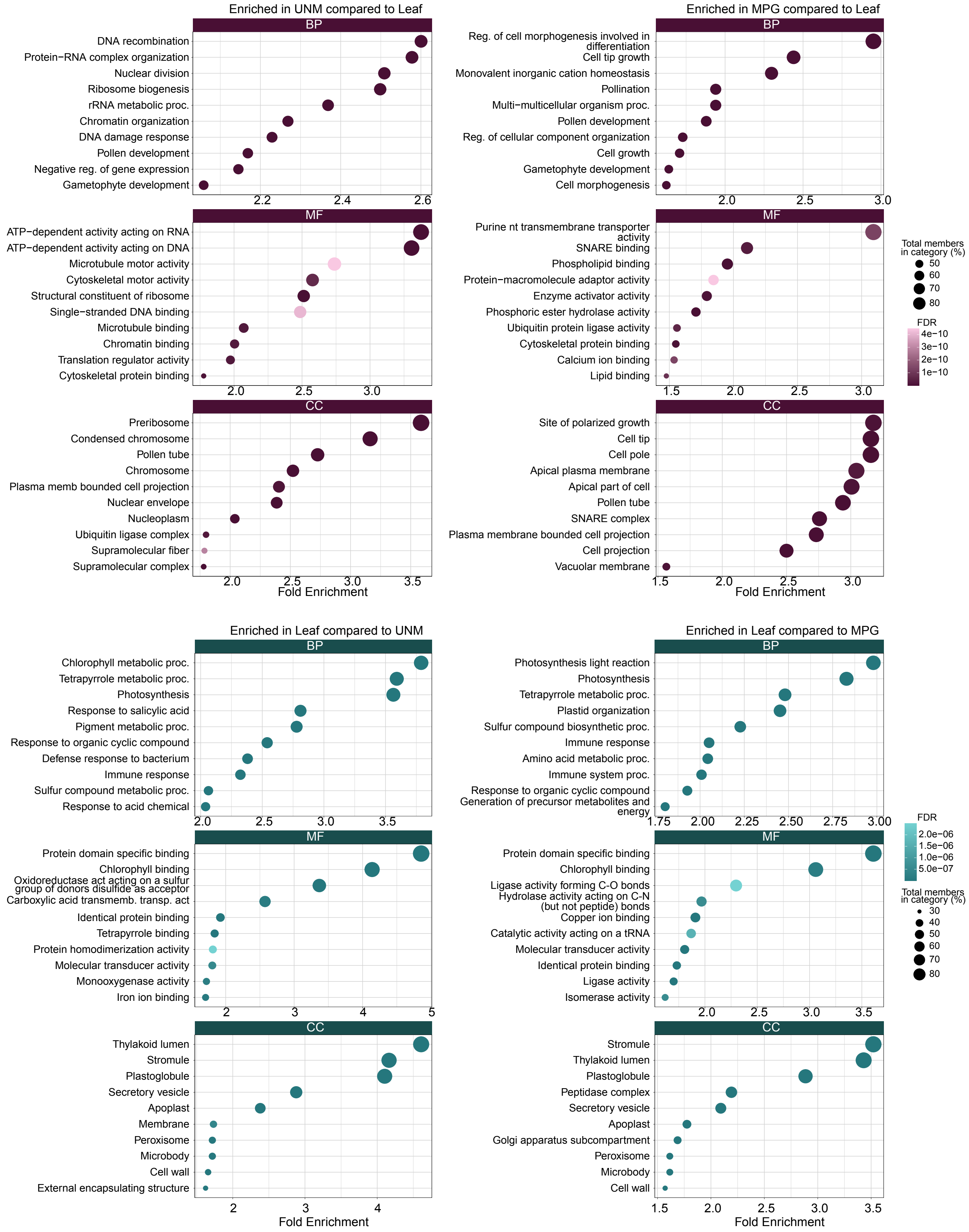

### Supplementary Figure 5

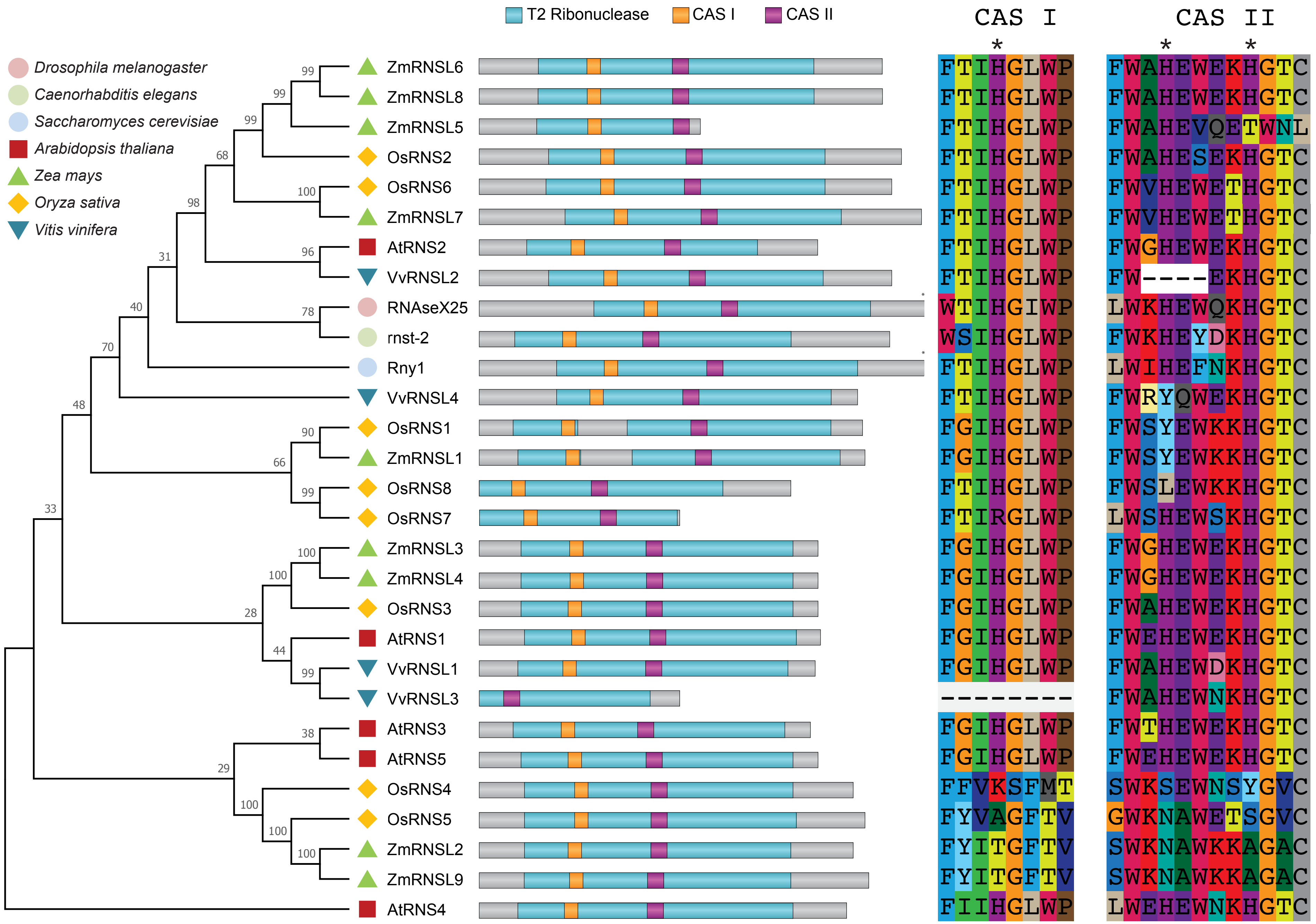

### Supplementary Figure 6

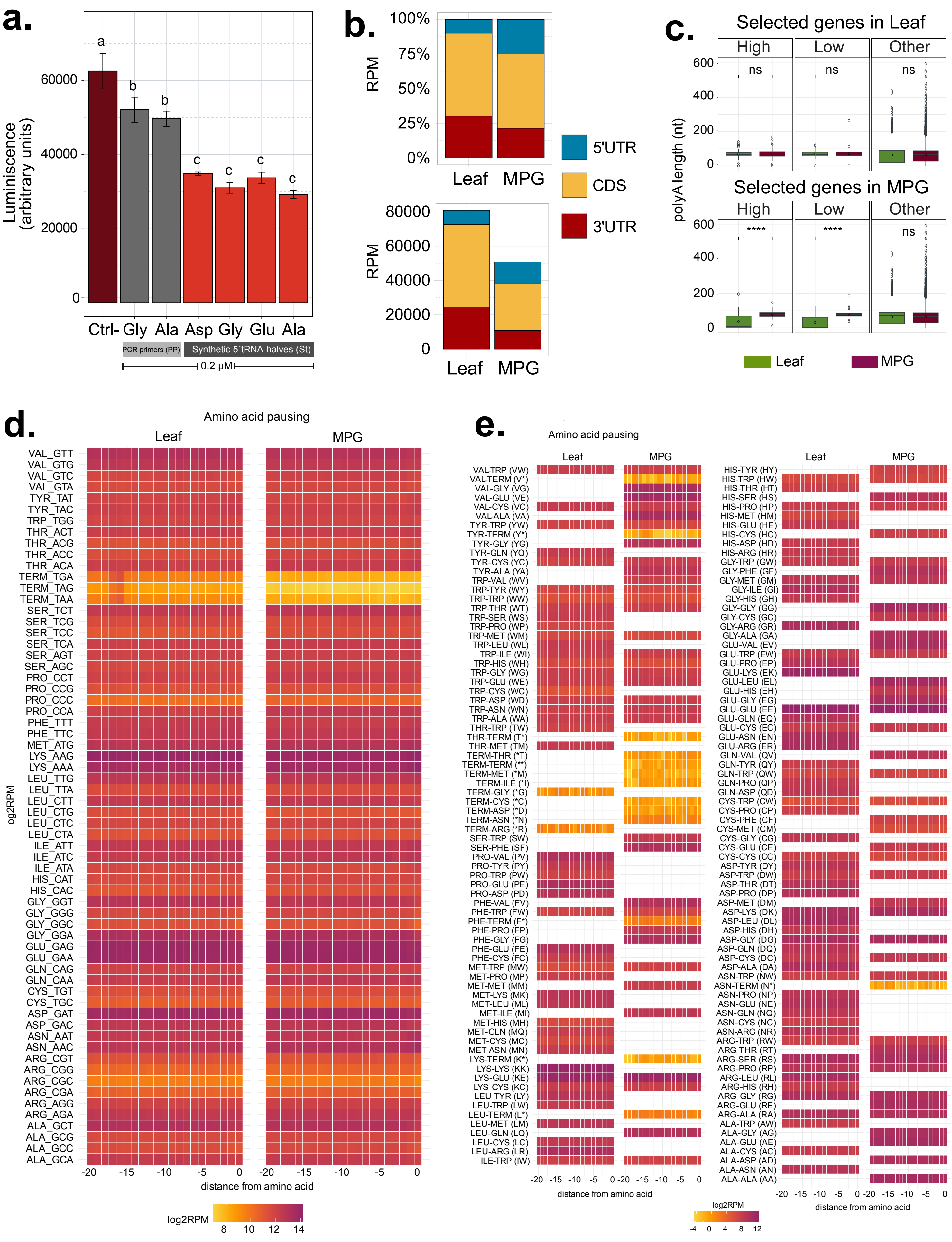

### Supplementary Figure 7

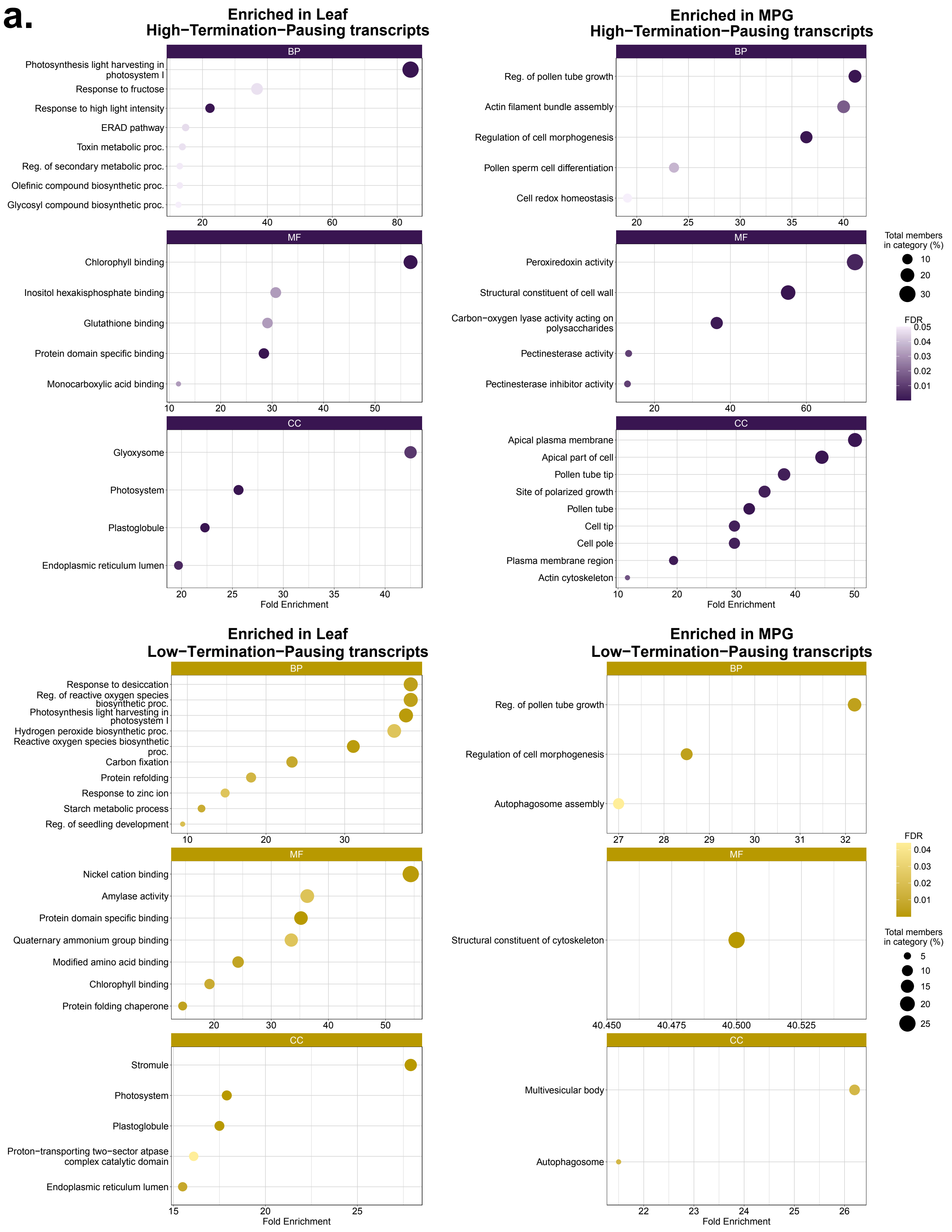

### Supplementary Figure 8

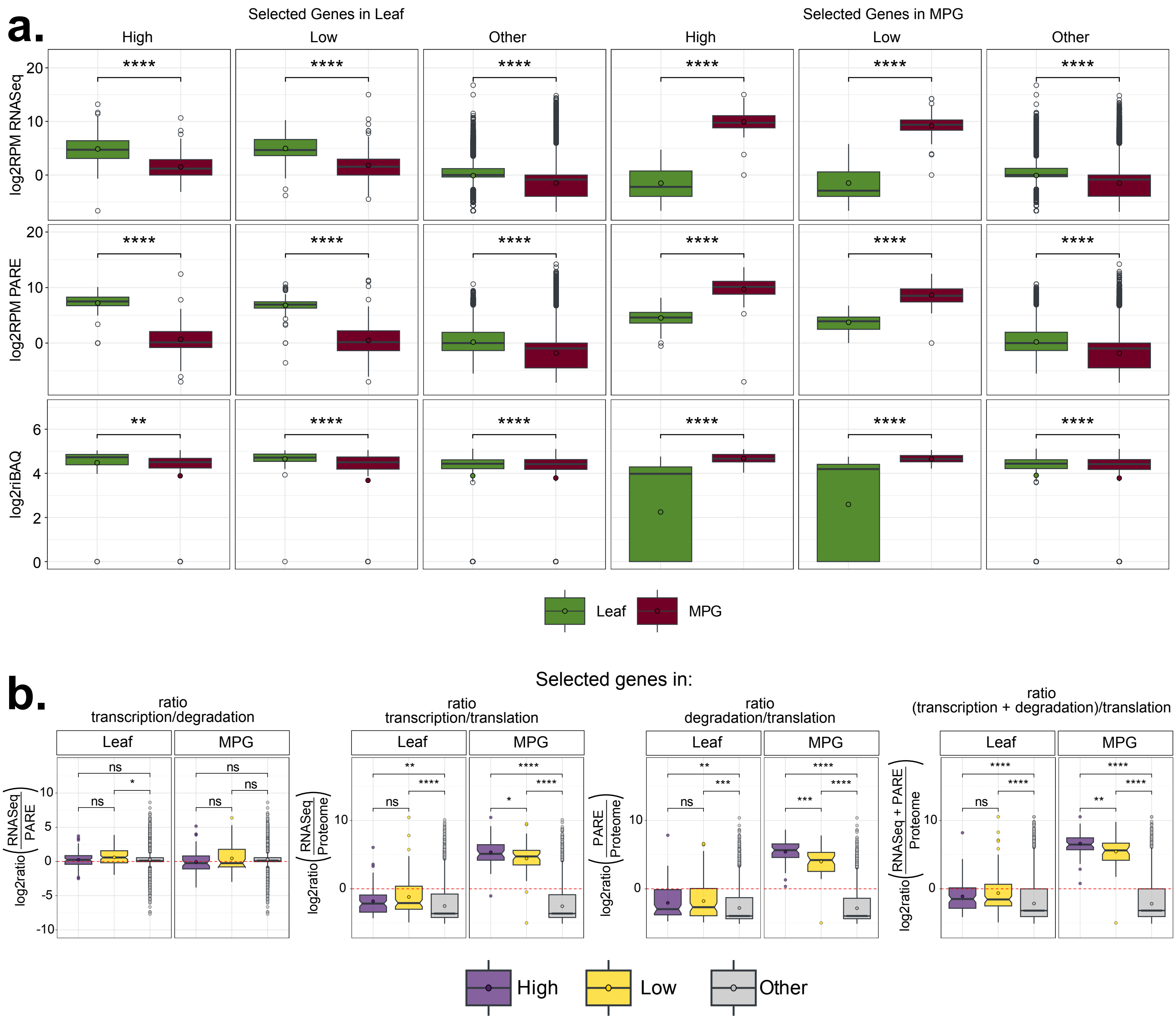

#### Supplementary Figure 9

**a.**

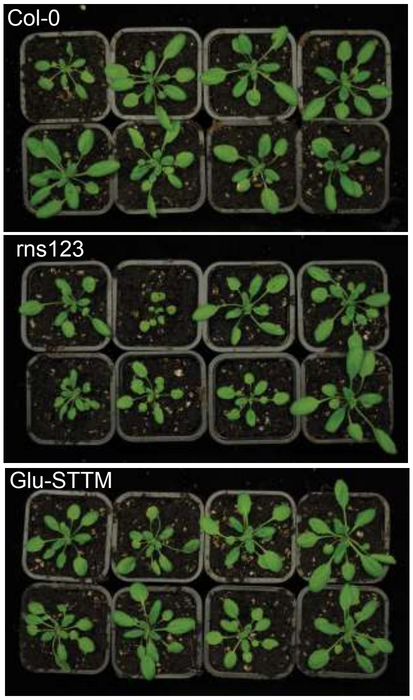


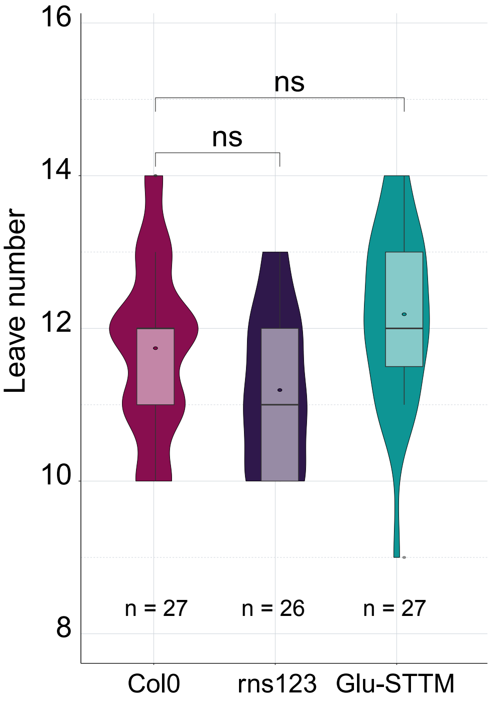


**b.**

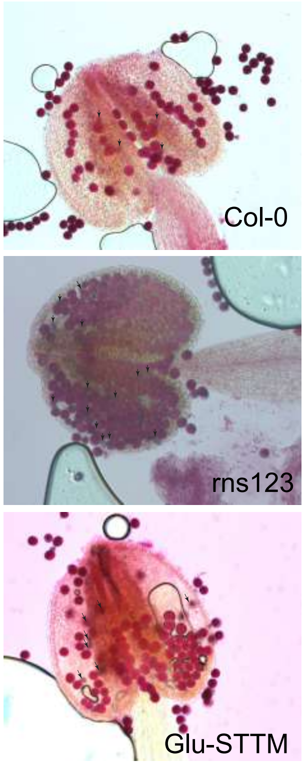


| | Viable pollen grains | Aborted pollen grains | % pollen viability |
|---|---|---|---|
| Col-0 | 98.5 | 953 | 15 |
| rns123 | 93.4 | 818 | 58 |
| Glu-STTM | 94.9 | 881 | 47 |

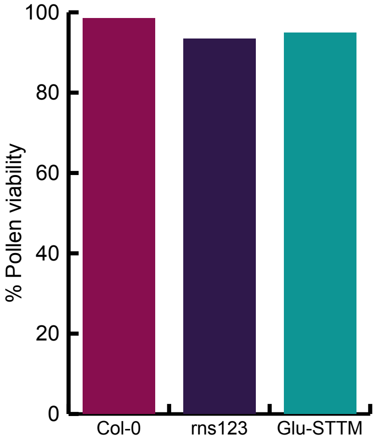
